# *Drosophila* Activin signaling promotes muscle growth through InR/dTORC1 dependent and independent processes

**DOI:** 10.1101/2020.03.23.003756

**Authors:** Myung-Jun Kim, Michael B. O’Connor

## Abstract

The Myostatin/Activin branch of the TGFβ superfamily acts as a negative regulator of mammalian skeletal muscle size, in part, through downregulation of insulin/IGF-1 signaling. Surprisingly, recent studies in *Drosophila* indicate that Activin signaling acts as a positive regulator of muscle size. Here we demonstrate that *Drosophila* Activin signaling promotes growth of the muscle cells along all three axes; width, length and thickness. In addition, Activin signaling positively regulates the InR/dTORC1 pathway and the level of MHC, an essential sarcomeric protein, via promoting the transcription of *Pdk1* and *Akt1*. Enhancing InR/dTORC1 signaling in the muscle of Activin pathway mutants restores MHC levels close to wild-type, but only increased the width of muscle cells. In contrast, hyperactivation of the Activin pathway increases the length of muscle cells even when MHC levels were lowered by suppression of dTORC1. Together, these results indicate that *Drosophila* Activin pathway regulates larval muscle geometry via promoting InR/dTORC1-dependent MHC production and the differential assembly of sarcomeric components into either pre-existing (width) or new (length) sarcomeric units depending on the balance of InR/dTORC1 and Activin signals.

## Introduction

Skeletal muscle accounts for a large portion of the body mass in various species including mammals (Gunn, 1989) and flying insects (Marden, 2000). It is essential not only for mobility but also for organismal energy balance and metabolism as it is a primary tissue for insulin-stimulated glucose consumption [reviewed in (Stump et al., 2006)]. Skeletal muscle is also proposed to be an endocrine organ that secretes a plethora of bioactive molecules, known as myokines, many of which depend on muscle contraction for production and secretion. Current evidence suggests that myokines exert substantial influence on the physiology and activity of their various target tissues [reviewed in (Pedersen and Febbraio, 2012)]. Therefore, achieving and maintaining an appropriate skeletal muscle mass and cellular function is likely to be essential for good health and quality of life.

Multiple signaling pathways are known to act in concert to achieve and maintain proper muscle mass [reviewed in (Piccirillo et al., 2014; Schiaffino et al., 2013)]. Among these, Myostatin (Mstn), a member of the TGF-β superfamily of growth and differentiation factors, has proven to be a prominent player. Loss-of-function mutations in *mstn* have been identified or induced in a large variety of mammals including mice, cattle, dogs, sheep and humans (Clop et al., 2006; Kambadur et al., 1997; McPherron et al., 1997; McPherron and Lee, 1997; Mosher et al., 2007; Schuelke et al., 2004) as well as birds and fish (Bhattacharya et al., 2019; Gao et al., 2016; McFarland et al., 2006). In all these species, loss of *mstn* results in larger skeletal muscles leading to the conclusion that Mstn is a negative regulator of muscle mass. Mechanistically, the increase in skeletal muscle mass caused by disruption of the *mstn* gene has, in some cases, been attributed to excess proliferation of muscle progenitors (hyperplasia), but is now thought to be primarily mediated by formation of a larger number of fibers, as well as to hypertrophy of each muscle fiber causing bigger cross-sectional area with no increase in myonuclei number (Amthor et al., 2009; McPherron et al., 1997; Lee et al., 2012; Sartori et al., 2009).

In addition to Mstn, Activins, have also been shown to negatively affect muscle mass (Chen et al., 2017; Chen et al., 2014). Mstn and Activins appear to synergize in suppressing muscle growth since co-inhibition of both factors resulted in a greater increases in muscle mass than those in which the activity of individual factors was inhibited (Chen et al., 2017) and expression of a dominant negative form of ActRIIB, a high affinity activin type 2 receptor for Mstn, Activins and several other ligands in the TGF-β/Activin subfamily (Lee and McPherron, 2001; Souza et al., 2008), leads to muscle hypertrophy (Lee and McPherron, 2001). Conversely, overexpression of these factors has been shown to promote the loss of muscle weight both in rats (Amirouche et al., 2009) and mice (Chen et al., 2014; Zimmers et al., 2002).

Several different muscle pathologies including cancer cachexia (Lokireddy et al., 2012; Loumaye et al., 2015; Marino et al., 2015), and muscle disuse due to paralytic injuries (Gustafsson et al., 2010; Reardon et al., 2001; Wehling et al., 2000) also shown a correlation between muscle loss and increased expression of Mstn or Activin. In the case of cancer cachexia and dystrophic muscle wasting, muscle lose can be partially reversed by inhibition of Mstn/Activin signaling (Bogdanovich et al., 2002; Chen et al., 2017; Zhou et al., 2010). Therefore, Mstn- and Activin-induced signaling pathways may provide potent therapeutic targets for the treatment of muscle atrophy in multiple clinical settings including age related sarcopenia (Bergen et al., 2015; White and LeBrasseur, 2014).

Although the vast majority of data strongly supports a negative role of Mstn/Activin signaling on muscle growth, there are some conflicting reports when manipulation of pathway activity is done at the R-Smad level. For example, Smad3 knockout mice exhibit smaller bodies and a ~10% reduction in the muscle fiber size (Tan et al., 2011) suggesting that Mstn/Activin regulation of muscle mass may be more complicated than is presently assumed.

A well-documented consequence of Mstn and Activin-induced signaling in the skeletal muscle is the inhibition of IGF-1/PI3K/AKT pathway. Specifically, overexpression of *mstn* via *in vivo* transfection in the adult muscle leads to attenuated phosphorylation of AKT, S6 and 4E-BP (Amirouche et al., 2009). Conversely, inhibition of Mstn and Activin by prodomain-derived antagonists or a soluble decoy receptor of ActRIIB leads to increased phosphorylation of mTOR and S6RP or increased phosphorylation of AKT and FOXO3a in skeletal muscle respectively (Chen et al., 2017; Zhou et al., 2010). These are all consistent with the idea that Mstn or Activin-induced signals lead to inhibition of the IGF-1/AKT/mTOR function the most important anabolic stimulus for muscle growth. (Egerman and Glass, 2014).

*Drosophila* skeletal muscles exhibit tremendous (around 50-fold) growth during larval stages (Piccirillo et al., 2014). This growth occurs through an increase in individual cell size without contribution from muscle stem cells (Demontis and Perrimon, 2009; Piccirillo et al., 2014). This process is mechanistically similar to mammalian muscle hypertrophy shown by *mstn* mutants that largely depends on growth of individual myofibers making *Drosophila* a good model for exploring the role of Activin signaling in regulating muscle fiber growth. In addition, the Activin signaling network is significantly simpler in *Drosophila* compared to vertebrates and consists of only three ligands, Activin beta (Actβ), Dawdle (Daw) and Myoglianin (Myo), a close homolog of vertebrate Myostatin, that signal through a single type I receptor, Baboon and a single R-Smad known as dSmad2 or Smox (reviewed in Upadhyay et al., 2017).

Intriguingly, we have recently shown that unlike vertebrates, the *Drosophila* Activin-like ligand Actβ is a positive regulator of larval muscle mass (Moss-Taylor et al., 2019). To distinguish whether this positive as opposed to negative growth function is a general feature of the entire Activin signaling network or represents an aberration due to loss of only one ligand, we analyzed muscle growth in *babo* and *dSmad2* null mutants which eliminate signaling of the entire pathway. We find that when the Activin signaling is completely compromised by loss of either *babo* or *dSmad2*, larval muscles are reduced in length, width and thickness similar to what we observe for *Actβ* loss alone. Hyperactivation of the Activin pathway through expression of constitutively activated Babo produces larger muscles in which length is disproportionally increased relative to width. We present evidence that Drosophila Activin signaling differentially controls larval muscle growth in three dimensions using both insulin dependent (width) and independent (length, thickness) pathways depending on the dose of the Activin signal.

## Materials and Methods

### *Drosophila* strains and husbandry

Fly lines were kept on standard cornmeal-yeast-agar medium at 25°C. For experiments involving *babo, dSmad2* and *daw* mutants, larvae were raised on yeast paste placed on apple juice-agar plates since the *babo, dSmad2* and *daw* mutants do not grow well on standard medium. The *w^1118^* strain was used as a wild-type (*wt*) control for *babo, dSmad2* and ligand mutants. *dSmad2^F4^, babo^fd4^, babo^df^, Actβ^ed80^, daw^1^, daw^11^*, *myo^1^, UAS-dSmad2* and *UAS-babo^CA^* (constitutively active) lines have been described previously (Brummel et al., 1999; Kim and O’Connor, 2014; Ting et al., 2007). *myo^CR2^* was generated by BestGene Inc by CRISPR-mediated mutagenesis using 5’-CTTCGACTATTCACCGCGCTATTA-3’ as a guide RNA. The resulting line contains a 1 bp deletion resulting in frameshift and stop prior to the ligand domain and is a putative null mutant. (Fig. S1E). The *Mef2-Gal4* (BL27390) line was used as a muscle driver throughout the study except in the RNA-seq analysis. To ensure that the results observed in this study are the consequence of muscle-specific acts of *Mef2-Gal4* driver, we repeated some of the key experiments using *Mhc-Gal4* (Demontis et al., 2014) driver which is considered to be more specific to skeletal muscle and obtained similar results (Fig. S3). Other stocks used are: *Pdk1^3^* (Rintelen et al., 2001), *Pdk1^33^* (Cheng et al., 2011), *UAS-dicer2* (BL24650), *UAS-S6kRNAi* (NIG 10539R-2), *UAS-S6KCA* (BL6914), *UAS-raptorRNAi* (BL31528-JF01087), *UAS-rictorRNAi* (BL31527-JF01086), *UAS-Pdk1* (Cheng et al., 2011), *UAS-Pdk1RNAi* (BL27725-JF02807), *UAS-Tor^DN^* (BL7013), *UAS-InR* (BL8284), *UAS-InR-RNAi* (VDRC 992), *UAS-Myc* (BL9675).

### Immunoblot analysis

Late foraging 3^rd^ instar larvae were used for immunoblots unless otherwise noted. Four to six larval body wall tissues containing muscle-epidermis complex were homogenized in 21 μl of RIPA buffer (Sigma, #R0278) supplemented with a cocktail of protease inhibitors (Complete mini, Roche) and incubated at 4°C for 40 min with agitation. After centrifugation, 13 μl of supernatant from each sample was transferred into a new tube, mixed with 7 μl of 3X loading buffer and denatured for 5 min at 95°C. Equal volumes from each sample were run on 4-12% Bis-Tris gel (Novex, #NP0322BOX) and transferred onto PVDF membrane (Millipore, #IPFL00010). The membranes were then blocked with Casein-containing buffer (Bio-Rad, #1610783) and incubated with primary antibody at 4°C overnight. The following primary antibodies were used: rabbit anti-AKT1 (1:1000, Cell Signaling Technology, #4691), rabbit anti-pAKT1 (1:1000, Cell Signaling Technology, #4054), rabbit anti-pS6K (1:250, Cell Signaling Technology, #9209), rabbit anti-pSmad2 (1:500, Cell Signaling Technology, #3108), mouse anti-α-Actn (1:50, DSHB, 2G3-3D7), anti-β-Tubulin (1:1000, DSHB, E7). Secondary antibodies were HRP-conjugated anti-rabbit or mouse IgG (1:10,000, Cell Signaling Technology, #7074 and #7076, respectively). Bands were visualized using Pierce ECL Western Blotting Substrate (Thermo Scientific, #32209) and band intensities were quantified using Image J (NIH) software. The quantification graphs are presented beneath the representative immunoblot images and the data are mean ± SEM from at least three independent samplings. The title of y-axis of each graph is ‘Relative protein level’ and is omitted for simplification.

### Muscle size assessment

Wandering larvae were rinsed in ddH_2_O and dissected in Ca^2+^-free HL3 as described previously (Kim and O’Connor, 2014). The larval fillets were then fixed in 3.7 % Paraformaldehyde solution (Electron Microscopy Sciences) for 15 min at RT. After washing in 1X PBS and permeabilization in 1X PBT (0.5% BSA+0.2% Triton X-100 in 1XPBS), the fillets were then incubated with Mouse α-Actn antibody (1:100, DSHB, 2G3-3D7) overnight at 4°C and Alexa555-conjugated secondary antibody (1:200, Molecular Probes) at RT for 2 hr.

Images were taken using Zeisss LSM 710 confocal microscope. From the α-Actn staining, the Z-disc numbers were counted using PeakFinder macro of Image J (NIH) followed by manual adjustment (Fig. S1C). The PeakFinder macro was also used in finding Z-disc intervals (Fig. S1C). The muscle width was measured using Image J software and normalized to wild-type or control. Muscle thickness was obtained from the orthogonal view of z-stack of 1 μm optical sections.

### Protein synthesis assay

Surface sensing of translation (SUnSET) method (Schmidt et al., 2009) was adopted to monitor protein synthesis capacity of the skeletal muscle with little modification. The SUnSET assay takes advantage of the fact that puromycin, a structural analog of aminoacyl-tRNA, can be incorporated into elongating polypeptide chains and can be immunologically detected. In the assay, 2-3 fillets of late foraging larvae were incubated in M3 insect medium (Sigma-Aldrich) supplemented with 2 mM of trehalose (Sigma-Aldrich), 15 μg/ml of puromycin (Sigma-Aldrich) and 10 μg/ml of insulin (Sigma-Aldrich) for 20 min at RT. The fillets were washed 3 times in 1X PBS and then sampled for immunoblotting. Anti-puromycin antibody (Millipore, #MABE343) was used at 1:10000. Representative images from triplicated assays are shown.

### qRT-PCR

Seven to ten larvae were dissected in 1X PBS to remove the internal organs. Total RNAs were prepared from the remaining muscle-epidermis complexes using TRIzol reagent (Invitrogen), followed by cleanup with RNAeasy Mini kit (Qiagen). The Superscript III first-strand synthesis kit (Invitrogen) was used to synthesize cDNA and qRT-PCR reactions were performed on LightCycler 480 (Roche) using SYBR green kit. Each sample was triplicated per reaction. *Rpl23* was used as a normalization control. The fold changes were calculated based on values obtained by 2^nd^ derivative maximum method. Data are mean ± SEM from at least three independent mRNA preparations.

### RNA sequencing (RNA-seq) analysis

Total RNAs were extracted using TRIzol reagent (Invitrogen) and further cleaned using RNAeasy kit (Qiagen) from thirty muscle-epidermis complexes of *wt* and *dSmad2* mutant as well as *tub-Gal80^ts^/+;Mhc-Gal4/+* and *tub-Gal80^ts^/+;Mhc-Gal4/babo^CA^* animals that were heat-shocked for 12 hr at 30°C for temporal expression of *babo^CA^*. Three μgs of total RNA per genotype were submitted to University of Minnesota Genomics Center (UMGC) for quality assessment and Illumina next-generation sequencing. In the UMGC, the integrity of RNAs was assessed using capillary electrophoresis (Agilent Bioanalyzer 2100). The sequencing libraries were constructed using TruSeq RNA preparation kit v2 after the mRNAs were enriched by oligo-dT-mediated purification. The libraries were then sequenced on a 100 bp paired-end run on the Illumina HiSeq 2500. Over 10 million reads were generated per library. The RNA-seq reads were mapped to *Drosophila* genome using TopHat. After mapping, the reads were assembled into transcripts using Cufflinks which generated fragments per kilo base of transcript per million mapped reads (FPKM) values. Gene differential expression test was performed using Cuffdiff. Finally, heat maps were drawn using R with ggplot2 package.

### Statistical Analysis

Statistical analyses were performed using Prism software (version 6.0, GraphPad Software). Data are presented as Mean ± SEM. One-way ANOVA followed by Dunnett’s test was used for comparisons among multiple groups and asterisks are used to denote the significance. Comparisons between two groups were performed by unpaired t-test and significances are denoted by pound signs. Graphs were drawn using either Prism or Exel software.

## Results

### Removal of the entire Activin signal pathway results in smaller larval muscles

To examine how muscle size is altered in Activin pathway mutants, we counted Z-discs from the larval skeletal muscle 6 of abdominal segment 2 stained with an α-Actinin (α-Actn) antibody. The Z-disc number is a proxy for sarcomere number and reflects the anterior-posterior length of the muscle cell. We also measured the lateral width of the muscle. We utilized heteroallelic combination of *baob^fd4^* and *babo^df^* (*babo^fd4/df^*), as well as *dSmad2^F4^*/Y as TGF-β/Activin pathway mutants in which the canonical signaling is completely abolished (Fig. S1A). The α-Actn antibody labels Z-discs with a similar intensity in wild-type and Activin pathway mutant muscles (Fig. 1A), consistent with the results from immunoblot analysis (Fig. 5A). Notably, however, the surface area of each muscle cell is smaller in the *babo* and *dSmad2* mutants (Fig. 1A). Quantitatively, the *babo* and *dSmad2* muscles exhibited ~35% reduction in Z-disc number (56.16 ± 1.39 for *wt* vs. 36.58 ± 0.78 for *babo* and 38.58 ± 0.71 for *dSmad2*) and ~25% decrease in muscle width (1.02 ± 0.02 for *wt* vs. 0.79 ± 0.02 for *babo* and 0.73 ± 0.03 for *dSmad2*) (Fig. 1B), demonstrating that the Activin signaling plays an essential role in new sarcomere additions leading to muscle lengthening as well as lateral expansion of sarcomeres adding to muscle width. In addition to the decrease in Z-disc number, the size of sarcomere measured by Z-disc interval is also decreased in *babo* but not in *dSmad2* mutant (Fig. S1D). Therefore, there appears to be additional effect on muscle length in *babo* mutant. Why the sarcomere size is affected by *babo* but not by *dSmad2* mutation is not clear however we have previously noted phenotypic differences between null alleles of these genes that arise through cross interactions of Babo and Mad (Peterson and O’Connor 2013; Kim and O’Connor, 2014). We also performed the same assay on abdominal muscle 7 of segment 2 (Fig. S1B) and muscle 6 and 7 of abdominal segment 3 (data not shown) and obtained the similar results. Therefore, it appears that the positive role of Activin signaling in muscle growth is not limited to certain muscle(s) but is likely general to all the muscles in the body wall (see also Taylor-Moss et al 2019). The effect of Activin signaling on muscle length and width is found to be cell-autonomous since expression of a *dSmad2* transgene using a muscle driver restored the decreased Z-disc number and muscle width of *dSmad2* mutant (Fig. 1C). To complete the assessment of muscle size along the 3 axes, we also measured muscle thickness and found that it is decreased by ~25% in *dSmad2* muscle (10.58 μm of *wt* vs. 7.81 μm of *dSmad2^F4^/Y;Mef2>+*, Fig. 1D and E). In addition, expression of a UAS-*dSmad2* transgene in the *dSmad2* muscle completely rescued muscle thickness (7.81 μm of *dSmad2^F4^/Y;Mef2>+* vs. 13.6 μm of *dSmad2^F4^/Y;Mef2>dSmad2*, Fig. 1E) indicating that Activin signaling cell-autonomously and positively regulates the thickness of muscle cell growth. We note that myonuclei number is not changed in *babo* and *dSmad2* mutants (Fig. S1F) indicating that the myoblast fusion occurs properly in these mutants.

**Fig. 1.**
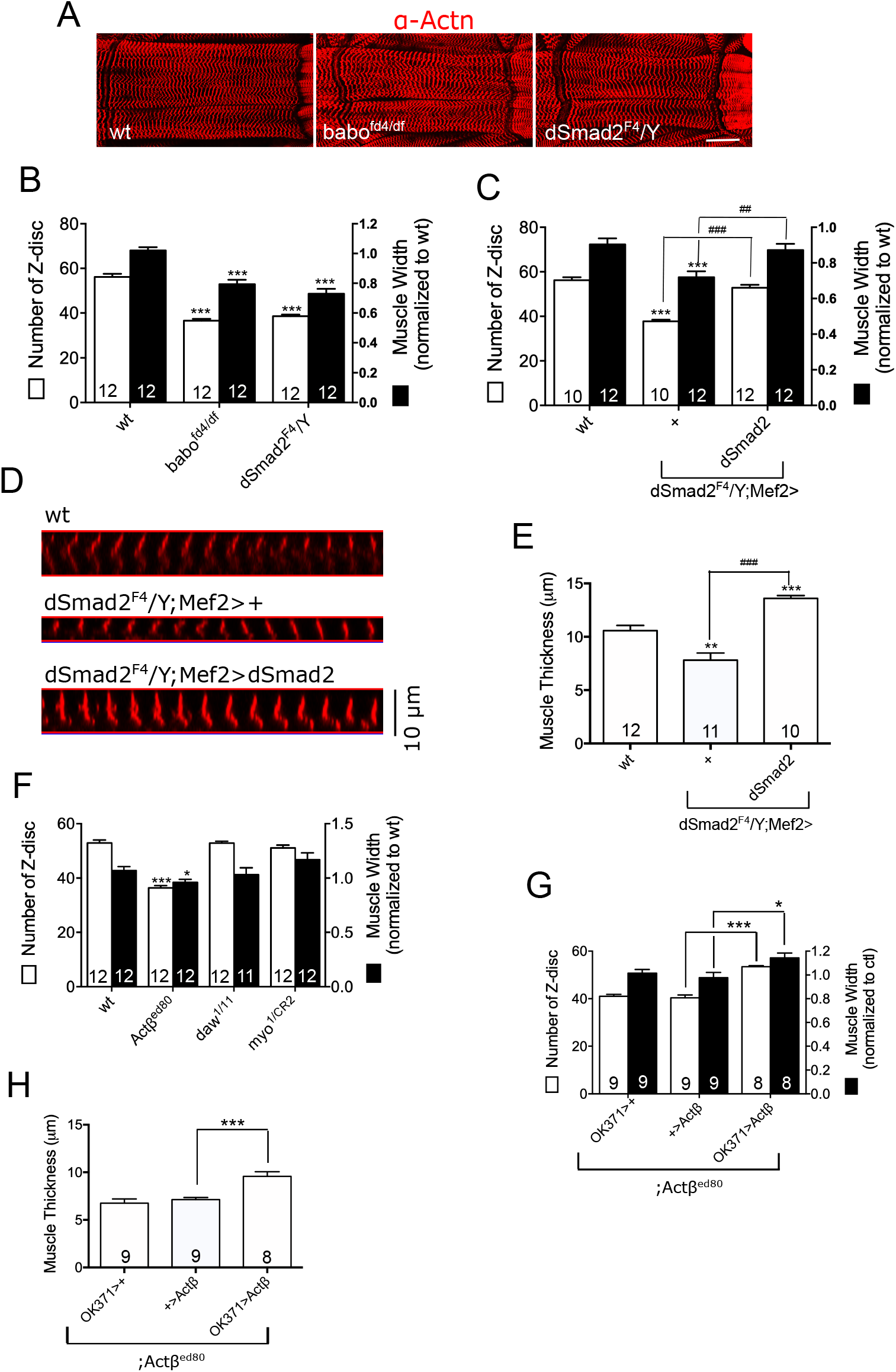
Activin signaling is necessary for proper muscle growth. (A) Representative images of muscle 6 of abdominal segment 2 of *wt* and Activin/TGF-β pathway mutants stained with α-Actn antibody. Scale bar equals 50 μm. (B) Assessment of muscle length by counting Z-discs and measurement of relative width of the muscle 6 of abdominal segment 2. Both the Z-disc number and muscle width are decreased in *babo* and *dSmad2* mutants. (C) Restoring Activin signaling in *dSmad2* muscle by expressing a wild-type *dSmad2* transgene rescues the reduced Z-disc number and muscle width. (D) Orthogonal optical z-stack sections illustrate muscle thickness (E) Expressing a wild-type *dSmad2* transgene rescues the reduced thickness of *dSmad2* muscle (F) Among ligands, only *Actβ* mutants display reduction in Z-disc number and muscle width. (G and H) Expression of *Actβ* in the motor neurons of *Actβ* mutant rescues the reduced Z-disc number (G) as well as thickness (H) of the muscle. Values are mean ± SEM. *p<0.05 and ***p<0.001 from one-way ANOVA followed by Dunnett’s test in which each genotype was compared to wt (B, C, E and F) or to *Actβ* /+; *Actβ^ed80^* control (G and H). Additionally, unpaired t-tests were performed in C and E as indicated by lines. ##p<0.01 and ###p<0.001 from unpaired t-test.

A similar reduction in the width and length of muscle cells is observed in *Actβ* mutants (Fig. 1F and (Moss-Taylor et al., 2019), but not in *myo* or *daw* mutants. Furthermore, overexpression of *Actβ* in motor neurons rescues the decreased Z-disc number, width (Fig. 1G) and thickness (Fig. 1H) of *Actβ* mutant suggesting that motor neuron-derived Actβ is the major Activin-like ligand that regulates Drosophila larval muscle growth. Interestingly, the skeletal muscle is disproportionately smaller when compared to other organs in *Actβ* mutants (Moss-Taylor et al., 2019). Together with the finding that Activin signaling regulates muscle size in a cell autonomous manner (Fig. 1C and E), these results indicate that the effect of Activin signaling on body size is primarily through regulation of muscle growth. Finally, was previously shown that myonuclei of *Actβ* mutant are smaller than those of wild-type (Moss-Taylor et al., 2019). To see if the reduction in myonculei size which is related to endoreplication is responsible for the smaller muscle phenotype of Activin pathway mutants, we overexpressed *Myc* in *dSmad2* mutant muscle to increase nuclar size. As illustrated in Fig. S4A and B, *Myc* overexpression greatly increased the myonuclei size, but failed to increase muscle surface area of *dSmad2* mutant. Furthermore, the ratio between myonuclei and muscle area is not significantly changed in *dSmad2* mutant (Fig. S4C). Taken together, these results demonstrate that Activin control of muscle growth does not solely involve myonuclei size and/or endoreplication.

### Influence of the Activin pathway on InR/dTOR signaling in larval skeletal muscle

In mammalian skeletal muscle, Mstn signaling is known to inhibit the IGF-1/PI3K/mTOR pathway (Amirouche et al., 2009; Chen et al., 2017; Zhou et al., 2010). To determine if the two pathways interact similarly in non-mammalian muscle, we investigated phosphorylation of AKT1 and S6K in *Drosophila* larval skeletal muscle-epidermis complexes of wild-type and Activin pathway mutants. The pAKT1 antibody used in this study detects phosphorylation of AKT1 at Ser505. This site corresponds to Ser473 of mammalian AKT and is phosphorylated by dTORC2 (Hietakangas and Cohen, 2007; Sarbassov et al., 2005; Yang et al., 2006). The pS6K antibody detects phosphorylation at Thr398 which corresponds to Thr389 of mammalian S6K and is phosphorylated by dTORC1 (Kockel et al., 2010; Lindquist et al., 2011; Sarbassov et al., 2005; Yang et al., 2006).

We first confirmed that phosphorylation at these sites is indeed dependent on InR activity in the larval skeletal muscle by overexpressing wild-type *InR* and *InR-RNAi* using *Mef2-Gal4* driver to increase or suppress the InR activity, respectively, in the skeletal muscle. As shown in Fig. 2A, phosphorylation of AKT1 is greatly reduced by *InR-RNAi* and increased by *InR* overexpression in the muscle-epidermis complexes, confirming that the phosphorylation level at Ser505 of AKT1 faithfully reflects the InR activity. The results also indicate that dTORC2 activation is one of the downstream events of InR activation since the Ser505 of AKT1 is exclusively phosphorylated by dTORC2 (Hietakangas and Cohen, 2007; Sarbassov et al., 2005). The phosphorylation at Thr398 of S6K is similarly regulated by InR activity, that is, decreased by *InR-RNAi* and increased by *InR* overexpression (Fig. 2A), suggesting that dTORC1 is also positively regulated by InR signaling in *Drosophila* larval skeletal muscle.

**Fig. 2.**
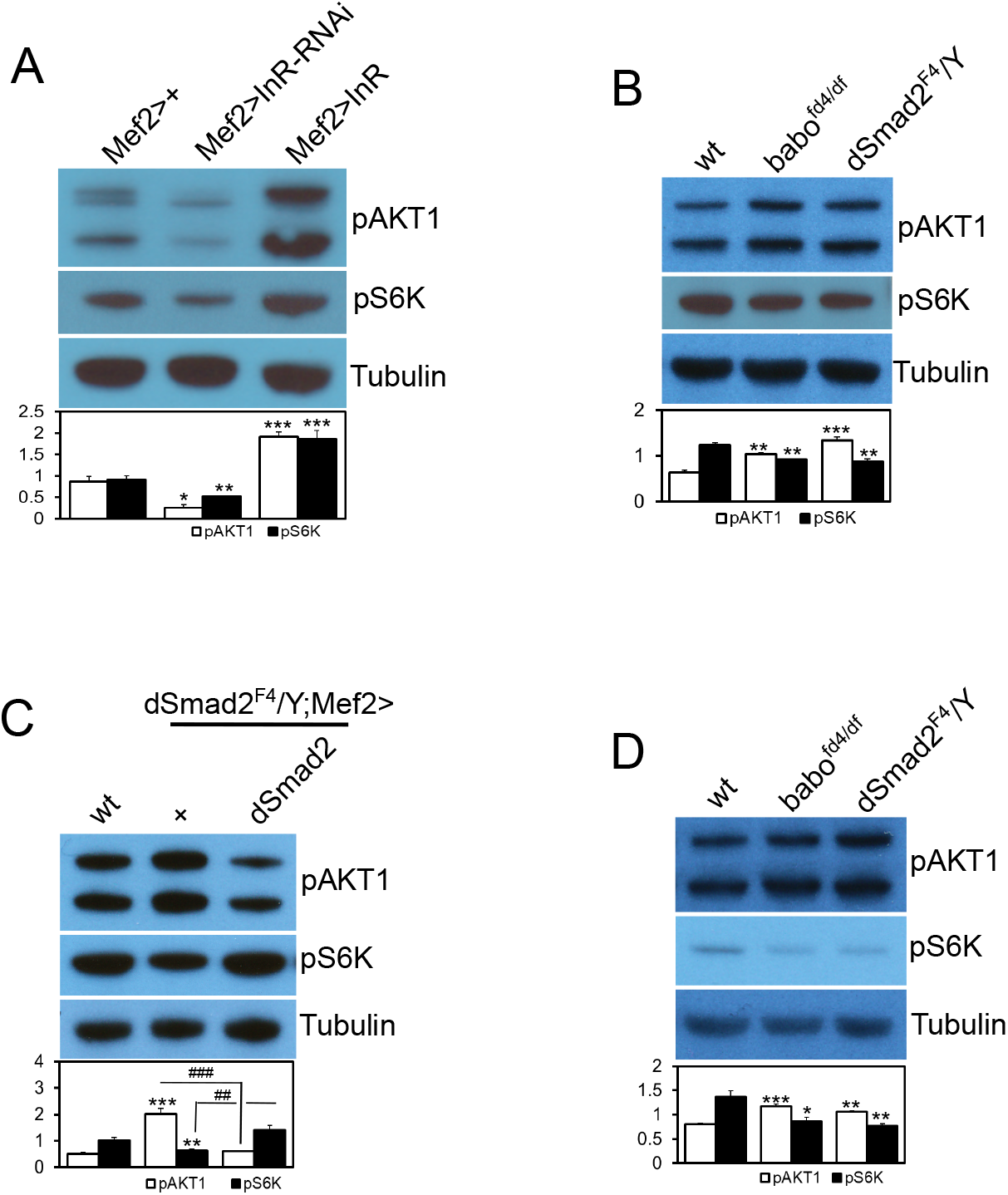
Activin pathway regulates the InR/dTOR signaling in the body wall. Representative immunoblot images and quantification of pAKT1 and pS6K. (A) Phosphorylation of AKT1 at S505 and S6K at T398 sites are down- and up-regulated by muscular expression of *InR-RNAi* and *InR*, respectively, suggesting that the InR signaling positively regulates the phosphorylation at these sites. (B) Phosphorylation of AKT1 is increased whereas the pS6K level is decreased in the larval body walls of *babo* and *dSmad2* mutants. (C) Resupply of Activin signaling in *dSmad2* muscle restores pAKT1 and pS6K levels close to wild type. (D) Wandering larvae display the same pattern of alteration in the AKT1 and S6K phosphorylation in the larval body wall as foraging larvae. Values are mean ± SEM. *p<0.05, **p<0.01 and ***p<0.001 from one-way ANOVA followed by Dunnett’s test in which each genotype was compared to *Mef2-Gal4/+* control (A) or *wt* (B, C and D). Additionally, unpaired t-tests were performed in C as indicated by lines. ##p<0.01 and ###p<0.001 from unpaired t-test.

We next examined the effect of loss of *babo* and *dSmad2* on phosphorylation of AKT1 and S6K. In mammalian skeletal muscle and myoblast culture, phosphorylation of AKT at Ser473 has been shown to be negatively regulated by Myostatin-induced Activin/TGF-β signaling (Lokireddy et al., 2011; Tan et al., 2011; Trendelenburg et al., 2009). We found increased phosphorylation at the corresponding site of *Drosophila* AKT1 (Ser505) in the *babo* and *dSmad2* mutants (Fig. 2B), which implies a negative effect of Activin/TGF-β signaling on AKT1 phosphorylation. The result also suggests that dTORC2 activity is elevated in *Drosophila* Activin/TGF-β pathway mutants. In contrast to AKT1, phosphorylation of S6K is mildly decreased in *babo* and *dSmad2* mutants (Fig. 2B), indicating a decreased dTORC1 activity. The influence of TGF-β/Activin signaling on phosphorylation of AKT1 and S6K appears to be muscle-specific and cell-autonomous in the muscle-epidermis complexes, since expression of a wild-type *dSmad2* transgene in the muscle of a *dSmad2* mutant resulted in a restoration of pAKT1 and pS6K levels toward those of wild-type (Fig. 2C). In addition to the continuously-feeding foraging larvae that are used for all the immunoblot analyses in this study, we also examined wandering larvae that have ceased feeding to determine if the alterations in pAKT1 and pS6K levels are dependent on feeding status. As in foraging stage, the wandering larvae of *babo* and *dSmad2* mutants also exhibited elevated pAKT1 and decreased pS6K levels (Fig. 2D). Taken together, these results indicate that the dTORC1 activity is down-regulated in Activin/TGF-β pathway mutants leading to a decreased phosphorylation of S6K (Thr398), while dTORC2 activity is upregulated resulting in an elevated phosphorylation of AKT1 (Ser505). Furthermore, the regulatory effects of Activin signaling on dTORC1 and dTORC2 activities appears to be independent of the feeding status.

### Negative feedback loop by S6K

Our findings suggest that the dTORC1 and dTORC2 activities are differentially affected by the loss of canonical Activin signaling while both of them are similarly regulated by InR activity (Fig. 2A). To determine why the phosphorylation states of AKT1 and S6K are changed in opposite directions, we examined if a negative feedback loop involving S6K played a role. It has been shown that mTORC1 negatively regulates the IGF-1/PI3K/AKT pathway by inducing S6K-mediated phosphorylation and degradation of insulin receptor substrate (IRS) (Harrington et al., 2004; Um et al., 2004). The inhibitory effect of S6K activation on AKT1 phosphorylation at Ser505 has also been demonstrated in *Drosophila* (Kockel et al., 2010; Sarbassov et al., 2005). Since the S6K activity is likely decreased in Activin pathway mutants judged by reduced phosphorylation at T398 (Fig. 2B) and the fact that there is lower protein synthesis capacity in these mutants (see below), it is possible that the elevated pAKT1 level in *babo* and *dSmad2* muscles is an indirect result of weakened inhibitory feedback of S6K on the InR-AKT1 axis. To test this possibility, we overexpressed an activated form of S6K (S6K^CA^) in *dSmad2* muscle to compensate for the decreased S6K activity and found that it decreases pAKT1 levels towards that of wild type (Fig. 3A). In addition, knockdown of *S6k* in wild-type muscle increased pAKT1 levels while overexpression of *S6k^CA^* decreased it (Fig. 3B). To further demonstrate the importance of negative feedback on the phosphorylation status of AKT1 at Ser505, we suppressed the dTORC1 activity by knocking-down *raptor*, a key component of dTORC1, and observed an increase in the pAKT1 level (Fig. 3C). Knocking-down *rictor*, a crucial component of dTORC2 complex, on the other hand, led to a decreased phosphorylation of pAKT1, further confirming that dTORC2 is the primary player in phosphorylating AKT1 at this site (Fig. 3C). Taken together, these results demonstrate that the elevated AKT1 phosphorylation in *babo* and *dSmad2* muscles is a consequence of decreased activity of dTORC1 and S6K. Finally, we investigated the protein synthesis capacity of wild-type and TGF-β/Activin pathway mutants which is known to be controlled by TORC1 and S6K activities. By adopting the SUnSET method (Schmidt et al., 2009), we find that protein synthesis capacity is reduced in the body wall tissue of *babo* and *dSmad2* mutants (Fig. 3D). Furthermore, the decreased capacity in protein synthesis was rescued by muscle specific expression of wild type *dSmad2* in *dSmad2* mutants (Fig. 3E). These results further demonstrate that the activities of dTORC1 and S6K, both key regulators of protein synthesis, are downregulated in the muscle of TGF-β/Activin pathway mutants.

**Fig. 3.**
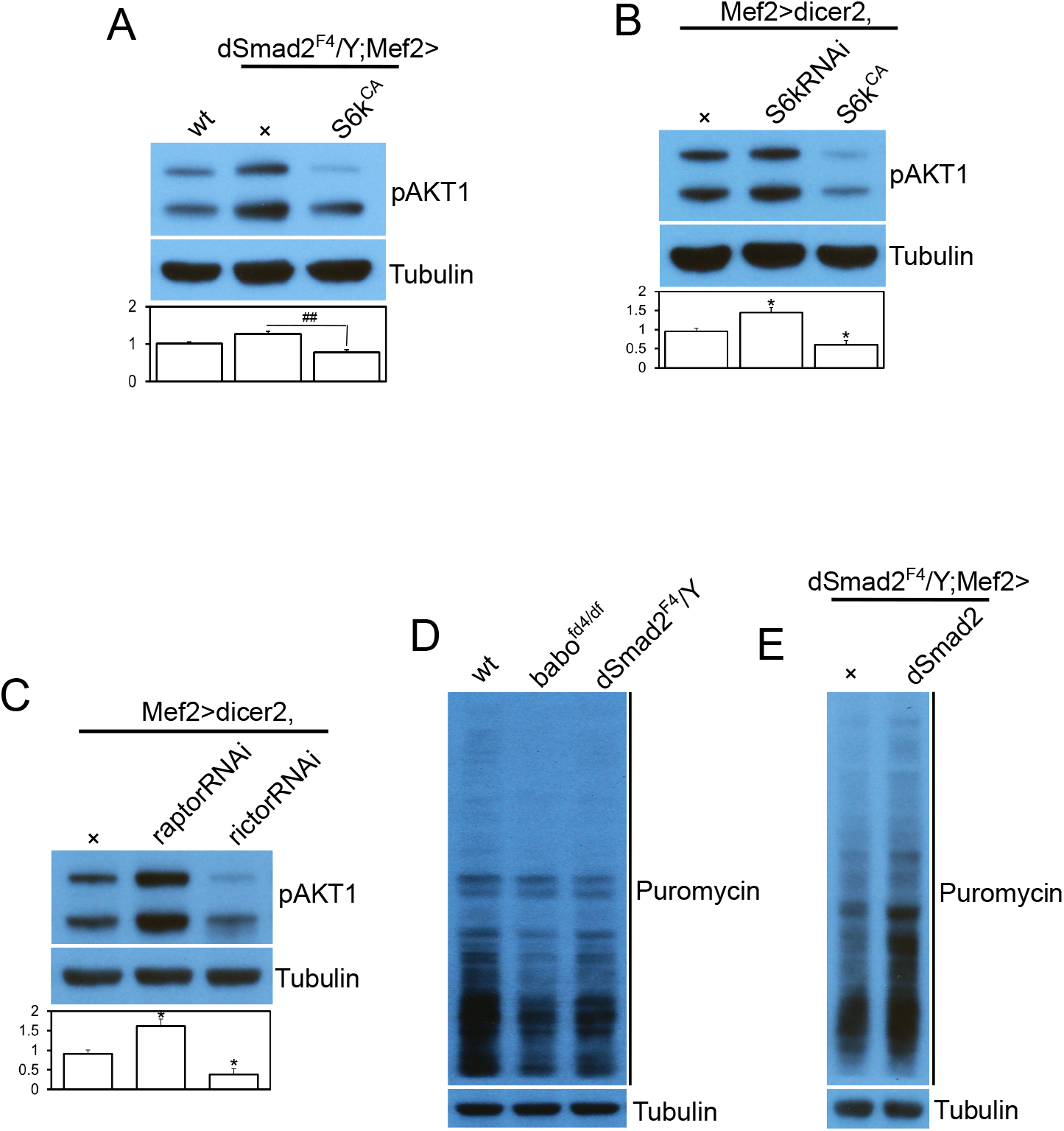
Increased phosphorylation of AKT1 is an indirect result of reduced inhibitory feedback by S6K. (A) Overexpressing a constitutively active form of *S6k* (*S6k^CA^*) suppresses the hyper phosphorylation of AKT1 in *dSmad2* mutant muscle. (B) Overexpression of *S6kRNAi* and *S6k^CA^* in wild-type muscle causes hyper-phosphorylation and hypo-phosphorylation of AKT1, respectively, indicating that the negative feedback loop from S6K to InR-AKT1 axis is functioning efficiently in larval body wall muscle. (C) Overexpression of *raptorRNAi* to inhibit dTORC1 activity results in an elevated pAKT1 level while *rictorRNAi* inhibits dTORC2 activity leading to decreased phosphorylation of AKT1. (D) The larval body wall tissue of Activin pathway mutants exhibits lower protein synthesis capacity assayed by SUnSET method. (E) Rescue of decreased protein synthesis capacity by muscle specific expression of *dSmad2* transgene. Values are mean ± SEM. *p<0.05 from one-way ANOVA followed by Dunnett’s test in which each genotype was compared to *wt* (A) or *UAS-dicer2/+; Mef2-Gal4/+* control (B and C). Additionally, an unpaired t-test was performed in A as indicated by lines. ##p<0.01 from unpaired t-test.

### Transcriptional regulation of InR/dTOR pathway components by Activin signaling

Since dSmad2 is a transcription factor, one possibility is that the *Drosophila* Activin pathway influences the InR/dTOR pathway via transcriptional control of one or more of its signal transduction component(s). To examine this issue, we performed RNA-seq using wildtype and *dSmad2* mutant as well as *Mhc-Gal4;tub-Gal80ts* control and *babo^CA^* gain-of-function-expressing samples. The FPKM values of genes encoding InR/dTOR signaling components (Fig. 4A) shows that transcripts of *Pdk1* and *Akt1* are significantly decreased while other InR pathway genes are unaffected in a *dSmad2* mutant suggesting a positive role of Activin/TGF-β pathway on the transcription of specific sets of InR/dTOR genes. In addition, temporal expression of activated Babo led to an increase in the transcripts of *Pdk1* and *Akt1* further demonstrating the positive role of Activin/TGF-β pathway (Fig. 4A). RNA-seq results of *Pdk1* and *Akt1* were then validated by qPCR analysis (Fig. 4B). Consistent with the findings from RNA-seq and qPCR analyses, the total protein level of AKT1 was found to be decreased in *babo* and *dSmad2* mutants (Fig. 4C), even though the pAKT1 level is elevated in these mutants (Fig. 2B). Taken together, these results indicate that the Activin pathway impinges on InR/dTOR pathway via controlling the transcription of Pdk1 and Akt.

**Fig. 4.**
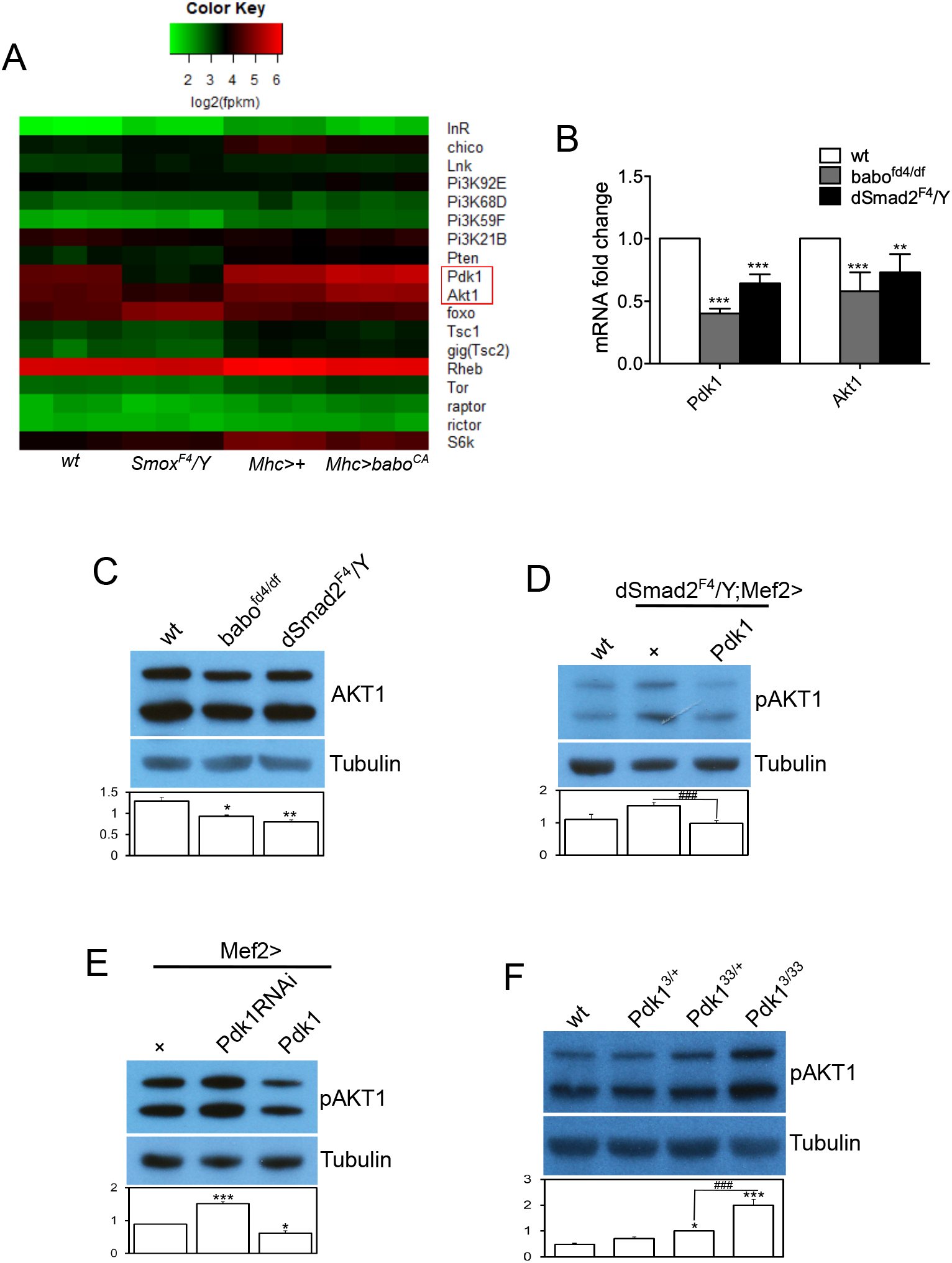
Activin signaling promotes the transcription of the InR/dTOR pathway components *Pdk1* and *Akt1*. (A) The heat map shows the effects of *dSmad2* loss and temporal over-expression of *babo^CA^* on the transcript levels of InR/dTOR pathway components. While most of the components are not affected, the expression of *Pdk1* and *Akt1*, highlighted by a red rectangle in the heat map, are downregulated by *dSmad2* mutation and upregulated by temporal expression of *babo^CA^*. (B) Verification of the RNA-seq results by qPCR. (C) Consistent with the RNA-seq and qPCR results, the total protein level of AKT1 is decreased in *babo* and *dSmad2* mutants. (D) Overexpression of *Pdk1* in *dSmad2* muscle restores the increased pAKT1 level towards that of wild type. (E) Expression of *Pdk1RNAi* in the skeletal muscle produces a similar phenotype in pAKT1 level as loss of Activin signaling while *Pdk1* overexpression suppresses the AKT1 phosphorylation. (F) A heteroallelic combination of *Pdk1* mutations causes hyper phosphorylation of AKT1. Values are mean ± SEM. *p<0.05, **p<0.01 and ***p<0.001 from one-way ANOVA followed by Dunnett’s test in which each genotype was compared to *wt* (B, C, D and F) or *Mef2-Gal4/+* control (E). Additionally, unpaired t-tests were performed in D and F as indicated by lines. ###p<0.001 from unpaired t-test.

In order to determine if the in Pdk1 and Akt transcription is responsible for any of the phenotypes exhibited by Activin pathway mutants, we sought to restore the expression level of *Pdk1* in the *dSmad2* muscle and examined pAKT1 (Ser505) level that inversely correlates with dTOCR1 and S6K activities (Fig. 3). As shown in Fig. 4D, overexpressing *Pdk1* in *dSmad2* muscle reduced the elevated pAKT1 level toward that of wild type, suggesting that the decrease in the expression of *Pdk1* is, at least partly, responsible for the elevated phosphorylation of AKT1. In a converse experiment, *Pdk1* was either knocked-down or overexpressed in otherwise wild-type muscles and the phosphorylation of AKT1 was examined. Consistent with the idea that *Pdk1* expression level negatively correlates with AKT1 phosphorylation, *Pdk1* knockdown increased pAKT1 levels while *Pdk1* overexpression decreased it (Fig. 4E). Since we showed above that the pAKT1 level also negatively correlates with S6K activity (Fig. 3B) and the S6K activity is enhanced upon phosphorylation at active site (Thr238) by PDK1 as inferred from mammalian results (Pullen et al., 1998), we propose that the effect of changes in *Pdk1* expression on pAKT1 phosphorylation is likely ascribed to the alterations in S6K activity. Finally, AKT1 phosphorylation is also found to be increased in *Pdk1* mutant (Fig. 3F) further supporting the idea of negative correlation between PDK1 activity and pAKT1 level. All together, these results indicate that one mechanism by which the Activin pathway influences the InR/dTOR signaling is via reduction in the transcription of several downstream signal transduction components can significantly affect the output of InR/dTOR signaling pathway even in the absence of a change in the ligand availability or activity and this appears to be the.

### Effect of Activin/TGF-β pathway on sarcomeric protein levels

In some cases, the Mstn /Activin pathway has been shown to negatively regulate MHC levels which in turn correlate with the change in muscle size in mammalian myoblast culture and skeletal muscle (Hulmi et al., 2013; Lokireddy et al., 2011). We examined if the Activin pathway similarly affects MHC levels in *Drosophila* muscle. In the immunoblot analysis using muscle-epidermis tissue, *babo* and *dSmad2* mutants exhibited reduced MHC abundance (Fig. 5A), even though the transcript level of *Mhc* is not changed (Fig. S2A). Furthermore, expressing a wild-type *dSmad2* transgene in *dSmad2* muscle rescued the decreased MHC level (Fig. 5C). In contrast, the total amount of α-Actn, another sarcomeric protein, is not altered in *babo* and *dSmad2* mutants (Fig. 5A) despite the increase in its transcript level (Fig. S2A). Therefore, it appears that the Activin pathway has variable effects on protein and transcript levels of different sarcomeric proteins.

**Fig. 5.**
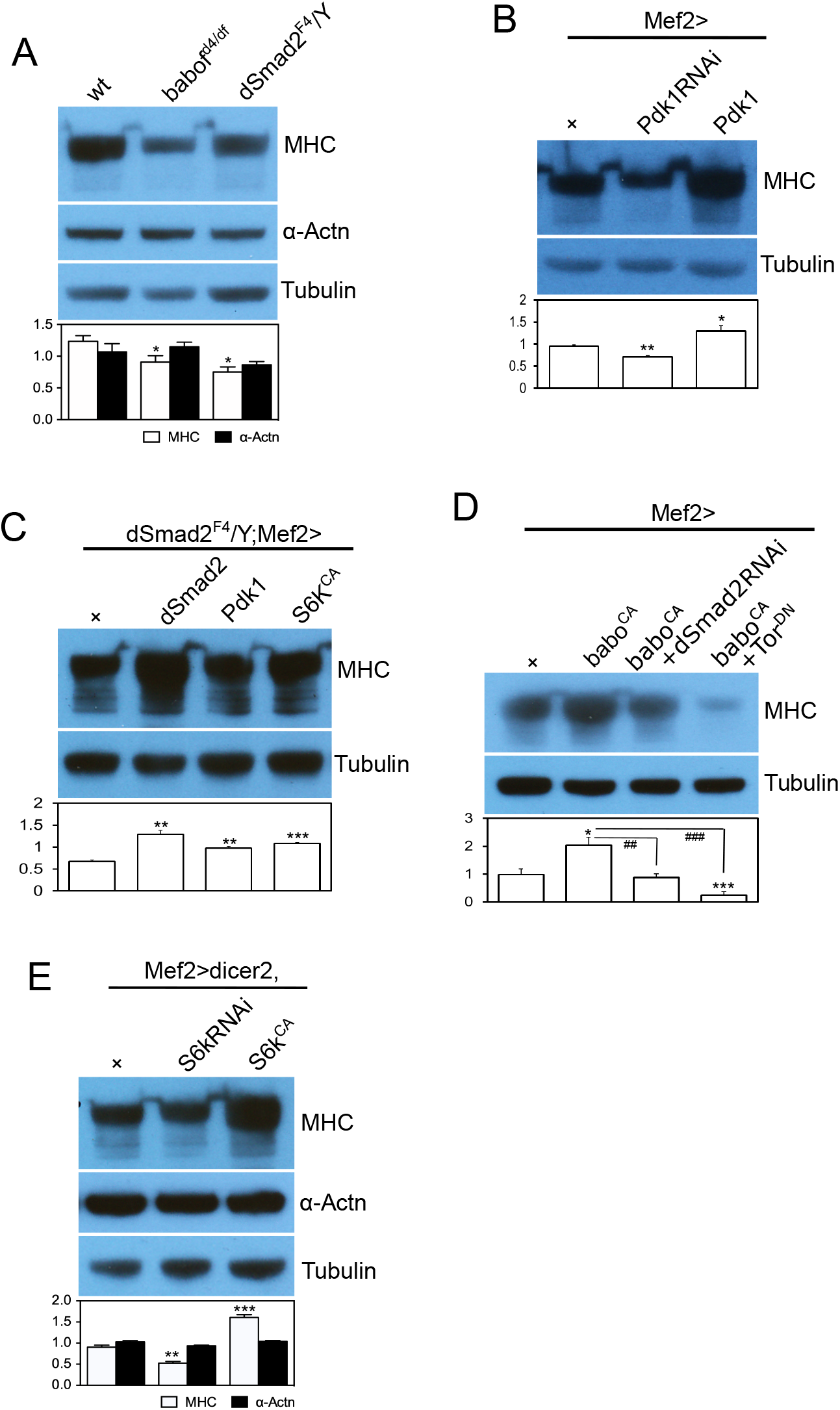
Activin signaling positively regulates MHC production through its effect on InR/dTOR1 activity. Representative immunoblot images and quantification of sarcomeric proteins. (A) The amount of MHC, a key sarcomeric protein, is decreased in the larval body walls of *babo* and *dSmad2* mutants while α-Actn, another sarcomeric protein that localizes to Z-discs, is not affected by these mutations. (B) Muscle expression of *Pdk1RNAi* decreases MHC abundance while expressing wild-type *Pdk1* in the muscle increases it, indicating a positive correlation between *Pdk1* expression level and the amount of MHC. (C) Expressing *dSmad2, Pdk1* and *S6k^CA^* transgenes in *dSmad2* mutant muscle rescues the decreased MHC level. (D) Expressing *babo^CA^* causes hyper-production of MHC which is suppressed by co-expression of *dSmad2RNAi* or *Tor^DN^*. (E) Expression of *S6kRNAi* and *S6k^CA^* in wild-type muscle reduces and increases the MHC level, respectively, without affecting the α-Actn level. Values are mean ± SEM. *p<0.05, **p<0.01 and ***p<0.001 from one-way ANOVA followed by Dunnett’s test in which each genotype was compared to *wt* (A), *Mef2-Gal4/+* (B and D), *UAS-dicer2/+;Mef2-Gal4/+* (E), or *Mef2-Gal4/+* control in *dSmad2* mutant background (C). Additionally, unpaired t-tests were performed in D as indicated by lines. ##p<0.01 and ###p<0.001 from unpaired t-test.

As shown above, a decrease in *Pdk1* expression gives rise to altered InR/dTOR signaling in *babo* and *dSmad2* mutants. Since the InR/dTOR signaling has a role in protein synthesis and degradation, we investigated if the decrease in *Pdk1* expression also contributes to the change in MHC level found in Activin pathway mutants. As illustrated in Fig. 5B, knockdown or overexpression of *Pdk1* resulted in decreased or increased levels of MHC, respectively, demonstrating a positive relationship between *Pdk1* expression and MHC protein levels. We then overexpressed *Pdk1* in *dSmad2* muscle and found that it rescues the decreased MHC level (Fig. 5C). From these results, we conclude that the decrease in *Pdk1* expression is, at least in part, responsible for the reduced MHC level shown by Activin pathway mutants.

Since we found that decreased *Pdk1* expression leads to diminished dTORC1 activity in Activin pathway mutants (Fig. 4D and E) we next investigated the contribution of dTORC1 to the regulation of MHC level by the Activin pathway. For this analysis, we expressed dominant negative *Tor (Tor^DN^*) together with activated *babo (babo^CA^*). Activated Babo alone increased the MHC production by 2-fold, which was suppressed by co-expression of *dSmad2RNAi* (Fig. 5D), meaning that the Babo^CA^ promoted MHC production primarily through canonical dSmad2-dependent signaling. Interestingly, co-expression of *Tor^DN^* resulted in an even stronger suppression of the hyper-MHC production induced by Babo^CA^ (Fig. 5D). Furthermore, a similar result was obtained by co-expression of *babo^CA^* and *raptorRNAi* (Fig. S2C) suggesting that dTORC1 activity mediates almost all of the effect of Babo^CA^ on the MHC level. Finally, we overexpressed *S6k^CA^* in *dSmad2* mutant muscle to assess the contribution of S6K, a key downstream effector of dTORC1, and found that it rescues the MHC level (Fig. 5C). Consistent with the essential role of S6K in regulating MHC production, expression of *S6kRNAi* and S6K^CA^ caused a significant decrease and increase in MHC levels, respectively (Fig. 5E). As in *babo* and *dSmad2* mutants, the α-Actn level is not affected by alterations in the S6K activity (Fig. 5E).

### Activin signaling promotes muscle growth through both InR/dTORC1 dependent and independent mechanisms

The findings that dTORC1 signaling, as well as MHC levels, are downregulated in Activin pathway mutants, led us to hypothesize that the reduction in MHC via the reduced dTORC1 signaling is primarily responsible for the decrease muscle growth observed in Activin pathway mutants. As shown above, the decreased MHC level of *dSmad2* muscle is rescued by overexpression of *Pdk1* and *S6k^CA^* (Fig. 5C). Because MHC is an essential building block of sarcomeres, we reasoned that overexpression of *Pdk1* or *S6k^CA^* would rescue the sarcomere number of *dSmad2* muscle. Surprisingly, however, overexpression of either of these transgenes failed to rescue the sarcomere number as assayed by counting the Z-discs in *dSmad2* muscle (Fig. 6A). Furthermore, the *S6k^CA^* even further decreased the sarcomere number from that of control *dSmad2* mutant muscle (Fig. 6A). Therefore, these results indicate that sarcomere formation can be decoupled from sarcomeric protein production and also suggests that the Activin pathway promotes sarcomere formation independently of its influence on InR/dTORC1 signaling and MHC production. Interestingly, overexpression of *Pdk1* or *S6k^CA^* increased the width (Fig. 6A) but not the thickness (Fig. 6B) of *dSmad2* muscle. From these results, we suggest that if MHC is over-produced in the absence of canonical Activin signaling, it is primarily used for lateral expansion of the muscle likely through addition to existing sarcomeres. To further test the role of Pdk1 in lateral growth of the muscle, we overexpressed *Pdk1-RNAi* to deplete PDK1 in otherwise wild-type muscle and found a decrease in muscle width while Z-disc number did not change (Fig. 6 C). We also altered S6K activity by overexpressing *S6kRNAi* or *S6k^CA^*. Despite profoundly affecting the MHC abundance (Fig. 5E), neither *S6kRNAi* nor *S6k^CA^*, had much effect on the Z-disc number (Fig. 6D), further demonstrating the lack of correlation between MHC production and formation new sarcomeres.

**Fig. 6.**
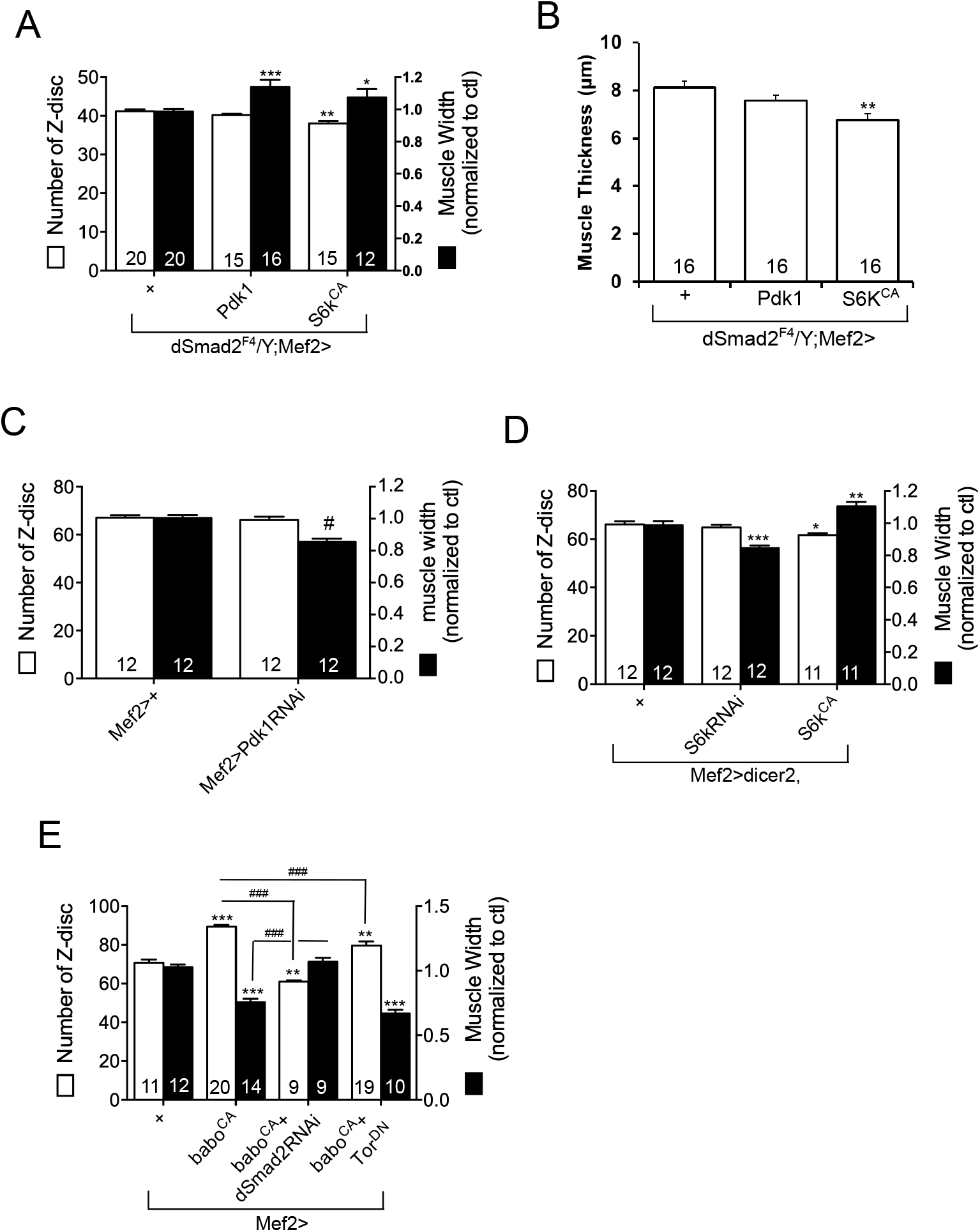
Z-disc number and muscle thickness are decoupled from MHC level. (A) Overexpression of *Pdk1* or *S6k^CA^* in *dSmad2* muscle rescues the muscle width but not the Z-disc number. (B) Overexpression of *Pdk1* or *S6k^CA^* failed to restore the reduced thickness of *dSmad2* muscle. (C) *Pdk1-RNAi* expression decreases the muscle width but not the Z-disc number. (D) Although the *S6kRNAi* and *S6k^CA^* profoundly affect the MHC level in larval body wall tissue (Fig. 5E), they have no or little effect on Z-disc number of the muscle. However, the muscle width is significantly reduced by *S6kRNAi* and increased by *S6k^CA^* expression. (E) Z-disc number and relative width of the muscle expressing *babo^CA^* alone and together with *dSmad2RNAi* or *Tor^DN^*. Values are mean ± SEM. *p<0.05, **p<0.01 and ***p<0.001 from oneway ANOVA followed by Dunnett’s test in which each genotype was compared to *Mef2-Gal4/+* control (E) or *Mef2-Gal4/+* control in *dSmad2* mutant background (A and B). Unpaired t-tests were performed in C and E (indicated by lines). #p<0.05, ###p<0.001 from unpaired t-test.

Finally, we counted Z-discs in muscles expressing *babo^CA^* together with *dSmad2RNAi* or *Tor^DN^*. Overexpressing *babo^CA^* alone causes an increase in the MHC level by 2-fold which was suppressed by co-expressed *dSmad2RNAi* or *Tor^DN^* (Fig. 5D). In the Z-disc counting assay, *babo^CA^*-expressing muscles exhibited, on average, 20 more sarcomeres than the *Mef-Gal4* controls (70.9 ± 1.43 for *Mef2>+* vs. 89.45 ± 0.91 for *Mef2>babo^CA^*; Fig. 6E). As expected, *dSmad2RNAi* completely blocked the increase in sarcomere number caused by *babo^CA^* overexpression and even further reduced the sarcomere number from that of control (89.45 ± 0.91 for *Mef2>babo^CA^* vs. 61 ± 0.72 for *Mef2>babo^CA^+dSmad2RNAi* vs. 70.9 ± 1.43 for *Mef2>+*; Fig. 6E). In contrast, the *Tor^DN^* only mildly suppressed the effect of *babo^CA^* so that the sarcomere number is still higher than that of control (89.45 ± 0.91 of *Mef2>babo^CA^* vs. 79.63 ± 2.13 for *Mef2>babo^CA^+Tor^DN^* vs. 70.9 ± 1.43 for *Mef2>+*; Fig. 6E). Considering that the *Mef2>babo^CA^+Tor^DN^* muscle likely has a higher level of Activin signaling but has a lower levels of MHC (Fig. 5D) than the *Mef2>babo^CA^+dSmad2RNAi* and *Mef2>+* control muscles, we conclude that the sarcomere number better correlates with the level of Activin signaling than with sarcomeric protein abundance. In contrast to the sarcomere number, muscle width is reduced by *babo^CA^* overexpression and *dSmad2RNAi* rescued it (Fig. 6E). We rationalize that the muscle width is smaller in *babo^CA^*-overexpressing muscle to accommodate the large increase in sarcomere number. In other words, sarcomeric subunits are assembled into new sarcomeres expanding the muscle length at the expense of widening of muscle through addition of sarcomeric proteins into existing Z-discs.

## Discussion

In this study, we assessed the effect of canonical *Drosophila* Activin signaling on InR/dTOR pathway activity and its relation to larval body-wall muscle growth. Our findings reveal an unexpected and striking difference in way that Activin signaling regulates muscle size in *Drosophila* larvae compared to mammals. In *Drosophila*, Activin signaling promotes muscle growth while in developing mammals it limits muscle mass. We find that the InR/TORC1 pathway is a core conserved target that mediates muscle size control in response to Activin, but the activity of the IGF-1/TORC1 pathway is regulated in the opposing direction compared to vertebrates. We also find that stimulation of InR/dTORC1 signaling in *Drosophila* in the absence of Activin leads to up-regulation of MHC which is incorporated into existing sarcomeres to increase their width. However, in the presence of Activin signaling width, length and thickness of muscle fibers are enhanced. The combinatorial effect of these sarcomeric assembly processes is the formation of larger larval body wall muscles with an increased volume.

### Differential modulation of IGF-1/TORC1 pathway accounts for the opposing effects of Activin signaling on mammalian verses *Drosophila* somatic muscles size

The present study demonstrates that the Activin pathway in *Drosophila* controls the output of the InR pathway by regulating the expression of PDK1 and AKT1, two downstream InR signal transduction components (Fig. 4). Since the steady-state levels of these two transcripts are lower in the body walls of *dSmad2* and *babo* mutants and are increased by expression of an activated Babo in muscle, it seems likely that these two genes are direct transcriptionally-regulated targets of dSmad2 and that Activin signaling boosts InR/dTORC1 activity resulting in enhanced S6K activity, higher general levels of protein synthesis, and increased levels of MHC. When reception of Activin signaling is compromised, then the opposite occurs. (Fig. 2 and 3). This mechanism is quite different from what has been proposed for how Mstn/Activin signaling impinges on the insulin/IGF-1 activity in mammals. In general, it has been reported that AKT phosphorylation is upregulated in the absence of Mstn/Activin signaling (Hitachi et al., 2014; Tan et al., 2011). Other points of intersection between the pathways have also been reported including several studies in mice suggesting that Mstn/Activin signaling suppresses expression of miRNAs that inhibit the *PTEN* translation (Goodman et al., 2013; Hitachi et al., 2014). This leads to lower levels of AKT phosphorylation, decreased mTORC1 activation and smaller muscles in the presence of Mstn/Activin signals.

The IGF-1/TORC1 pathway is only one point of intersection from which to understand muscle size control. In general, muscle homeostasis is thought to be regulated by balancing the activities of protein synthesis and degradation pathways (Bonaldo and Sandri, 2013; Schiaffino and Mammucari, 2011). The IGF-1/TORC1 signaling clearly interfaces with both these modes of protein homeostasis control, however, in most cases it is not clear whether Smad directly regulates expression of specific components in either the synthesis or degradation pathways or whether most of its affects can be attributed to regulation of IGF-1/TORC1.

### IGF-1/TORC1 negative feedback and muscle homeostasis

It is well documented in mammals that there is a negative feedback loop formed by S6K toward IRS which profoundly diminishes the efficacy of signaling from insulin/IGF-1 to PI3K-AKT axis (Harrington et al., 2004; Shah et al., 2004; Zhang et al., 2008). A similar negative feedback loop has also been demonstrated in *Drosophila* cell culture (Sarbassov et al., 2005) and wing imaginal discs (Kockel et al., 2010), but has never been studied in skeletal muscle. Here we demonstrate that this negative feedback loop does indeed work efficiently in *Drosophila* skeletal muscle. As the inhibitory feedback has a profound effect on insulin responsiveness, it will be interesting to determine how the peripheral tissues in Activin pathway mutants react to *Drosophila* insulin-like peptides. Together with the fact that the absolute expression level of PDK1 and AKT1 are down-regulated in the Activin pathway mutants, there may not be a resultant enhancement of the responsiveness. Consistent with the idea, we previously reported that the *dSmad2* as well as *daw* mutants display increased hemolymph sugar levels (Ghosh and O’Connor, 2014), suggesting an impairment in the regulation of blood sugar level. In addition, it might also indicate that the decrease in PDK1 and AKT1 expression overrides the effect of relieved negative feedback. Further study is required to unveil how these competing effects are summed by tissues to determine their responsiveness to insulin or other growth factors.

### Which TGF β superfamily members control muscle size

In mammals, the TGFβ superfamily consists of at least 30 ligands which are broadly classified into the BMPs that transduce signals through Smads1,5,8 and members of the TGFβ/Activin subgroup, including Mstn, that signal through Smads2 and 3 (Sartori et al., 2014). The *Drosophila* system is much simpler with only 6 clear family members, three of which are classified as BMPs and three that belong to the Activin subgroup including Myoglianin, the homolog of vertebrate Mstn (Upadhyay et al., 2017). In vertebrates, the full complement of ligands that participate in muscle size control in not known. Mstn is by far the best-studied family member in terms of post-myogenic muscle growth control, however studies employing overexpression in mice of various types of ligand binding proteins with diverse specificity for the Activin family members produced more extreme muscle hypertrophy than the *mstn* knockout alone, implicating that other Activin/TGFβ factors likely contribute to muscle growth control (Chen et al., 2015; Chen et al., 2017; Chen et al., 2014; Lee and McPherron, 2001; Lee et al., 2005; Winbanks et al., 2012).

While the BMP arm of the superfamily has not received as much attention, overexpression of BMP-7 or its activated type I receptor in muscles resulted in enhanced Smad1,5,8 phosphorylation and hypertrophic muscle growth (Sartori et al., 2013; Stantzou et al., 2017; Winbanks et al., 2013). Intriguingly, this appears to be accomplished, in part, through mTORC1 activation, increased protein synthesis, and reduced protein turnover, very similar to what we find for the Activin pathway in *Drosophila*. At present, no specific studies addressing the role of BMPs in muscle growth control have been reported in *Drosophila*, although it is worth noting that the BMP-7 homolog, Gbb, is expressed in larval muscle and strongly affects NMJ size and function (McCabe et al., 2003).

In terms of three *Drosophila* Activin-like ligands, muscle size regulation appears to be primarily accomplished by motoneuron delivery of Actβ to the muscle during larval growth (Moss-Taylor et al., 2019; this report). Loss of Actβ results in reduction of larval muscle size to a similar extent as we find here for Babo and dSmad2 loss, while genetic null mutations in either *myo* or *daw*, the other two activin-like ligands, produces no change in muscle size (Fig. 1D). Furthermore, loss of Actβ also results in similar electrophysiological defects at the NMJ as found for *babo* and *dSmad2*, while loss of Daw or Myo have little effect (Kim and O’Connor, 2014). These data all support Actβ as the primary *Drosophila* TGFβ-like ligand involved in muscle size control.

It is surprising that we find no effect of *myo* loss on muscle size since it is a clear homolog of vertebrate Mstn. Furthermore, it has been reported that RNAi knockdown of *myo* in muscles does result in a size increase, (Augustin et al., 2017). At present, we do not know why the RNAi results are different from the genetic null data. Discrepancies between the tissuespecific RNAi knockdown and genetic null phenotypes have been reported quite frequently in both *Drosophila* and vertebrates. One possible explanation is the activation of compensatory pathways in the null mutant that are not elicited by tissue specific knockdown methods (Di Cara and King-Jones, 2016; El-Brolosy et al., 2019; El-Brolosy and Stainier, 2017; Gibbens et al., 2011).

A major unanswered question concerns how the formation of new sarcomeres vs expansion of existing sarcomeres is differentially regulated. Interestingly, it has recently been reported that overexpression of the PR isoform of Zasp52, a *Drosophila* ALP/Enigma family protein, results in an increased myofibril diameter in adult indirect flight muscle (González-Morales et al., 2019), indicating that Zasp52-PR promotes the lateral expansion of sarcomeres. The result, however, does not give an insight into whether Zasp52-PR is also involved in serial addition of sarcomeres that will cause lengthening of the muscle, since the observation is limited to adult indirect flight muscle whose length is pre-determined during the pupal stage. In addition to Zasp52 isoforms, the effects of other Zasps were also investigated in the study and it was found that overexpression of Zasp66 or Zasp67 increases the population of myofibrils having either smaller or larger diameters, indicating that these two Zasps have dual roles in determining myofibril diameter. Interestingly, we found that Zasp66 expression is decreased in larval skeletal muscle of *babo* and *dSmad2* mutants (Fig. S2B) which may be contributing to the muscle size phenotypes and warrants further study.

Another consideration should be the influence of mechanical force generated at the muscle-tendon junction. It has been shown that attachment of flight muscle to the tendon during metamorphosis generates tension at the muscle-tendon junction which, in turn, triggers addition of sarcomeres in a head-to-tail manner during metamorphosis (Weitkunat et al., 2014). It is not clear if mechanical force exerts a similar effect in larval skeletal muscle since the larval exoskeleton is composed of soft cuticle that keeps changing in length as larvae move, a very different situation from that of pupae and adults in which tendon/muscles are attached to stiff cuticle. Additional study will be required to resolve this issue.

### Evolutionary aspects of Activin signaling and muscle size control

Why mammals and *Drosophila* are wired in opposite directions in terms of Activin signaling’s influence on IGF-1/TORC1 activity and muscle size control is unclear. The Mstn/ Activin branch of the TGF-β family is very ancient and is present in some pre-Bilateria groups including several cnidarian species (Watanabe et al., 2014), but functional studies of its role in muscle size control in these animals are lacking. Interestingly, there are two reports that examine Mstn function in crustaceans, one using the giant prawn *Macrobrachium rosenbergii* (Easwvaran et al., 2019) and the other the penaeid shrimp *Penaeus monodon* (De Santis et al., 2011). Despite both species being members of the Malacostraca class of Crustacea, these studies reached opposite conclusions suggesting that Mstn activity had an apparent positive role in body growth in *Penaeus* (De Santis et al., 2011), while in *Macrobrachium* it had a negative effect on muscle size (Easwvaran et al., 2019). Clearly, many additional phyla, classes and species of animals need to be examined to more fully understand how and why these contrasting roles in muscle size control have evolved.

In summary, we envision a two-step mechanism for how Activin controls *Drosophila* muscle fiber growth (Fig. 7). On one hand, it regulates production of muscle fiber structural subunits such as MHC through enhancement of *Akt1* and *Pdk1* transcript levels. These subunits can be assembled laterally to existing sarcomeres to build muscle fiber width. In a second step, we propose that it either positively regulates the production of a sarcomere initiation factor(s) or blocks the destruction of such components leading to length wise addition of new sarcomeric units and also to thickening of the muscle. It is important to recognize that although some physiological defects have been noted in muscles of *Actβ, dSmad2* and *babo* mutants (Kim and O’Connor, 2014), these do not cause visibly noticeable changes in either larval motion or feeding behaviors. Therefore, just as in the case of mammalian Mstn signaling, *Drosophila* Activin signaling is not essential for earlier steps of myoblast fusion and muscle differentiation and function, rather it appears to be a mechanism for fine-tuning muscle fiber growth.

**Fig. 7.**
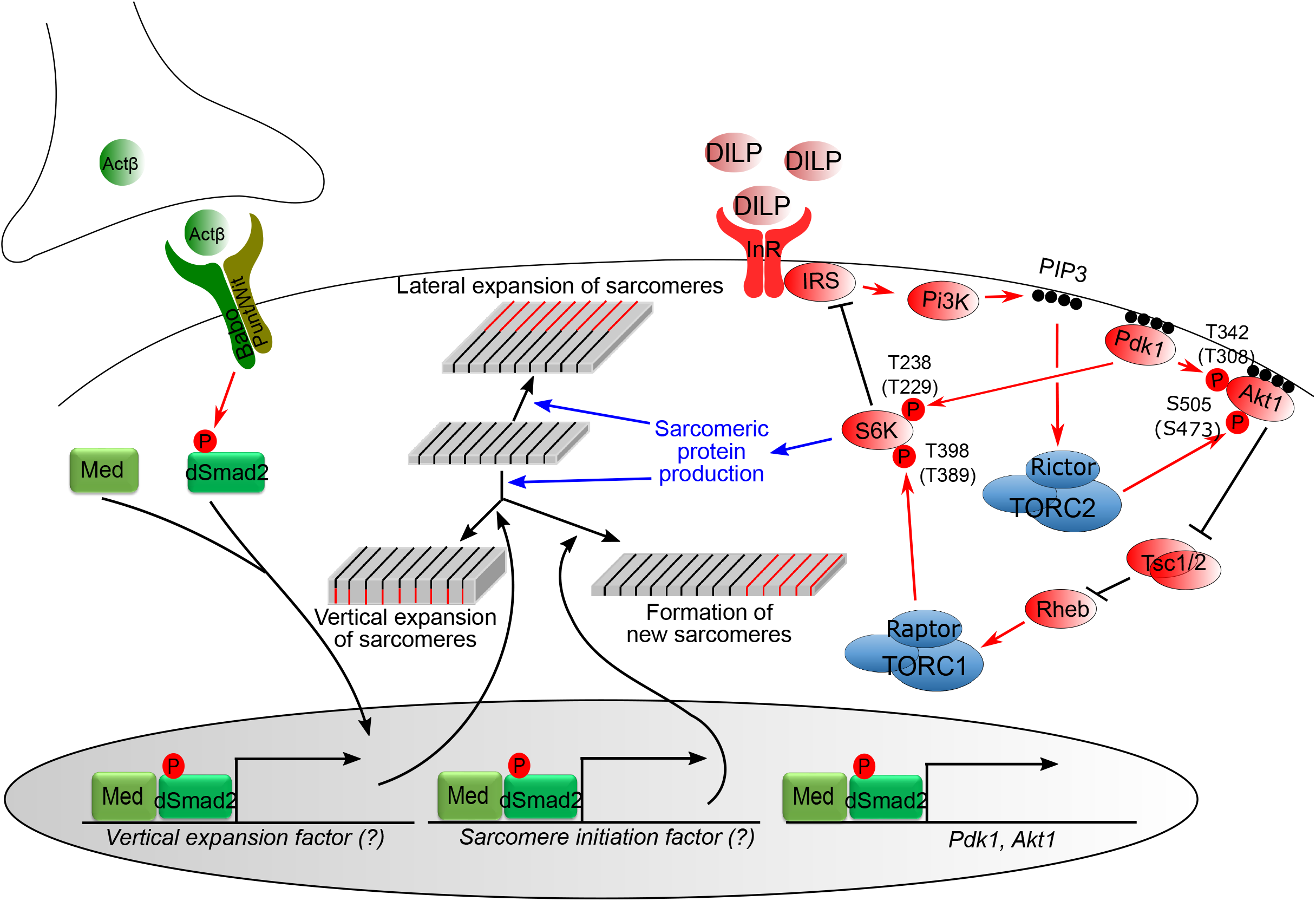
Control of InR/dTOR signaling network and muscle growth by Activin signaling pathway. Activin signaling positively regulates InR/dTOR signaling by promoting the transcription of *Pdk1* and *Akt1*. Activation of InR increases PI3K-dependent PIP3 generation leading to increased activity of PDK1. PDK1 then phosphorylates AKT1 at Thr342 and S6K at Thr238. The PI3K-generated PIP3 is also necessary for activation of the dTORC2 complex which phosphorylates Ser505 of AKT1. When phosphorylated at Thr342 and Ser505 sites, AKT1 initiates a cascade of inhibition leading to activation of dTOCR1 complex that phosphorylates S6K at Thr398. The sequential phosphorylations at Thr398 and Thr238 sites fully activates S6K. The activated S6K then promotes the production of certain sarcomeric proteins as well as inhibits signal transduction from InR to PI3K. The InR/dTORC1 signaling increases the steay-state level of MHC which is preferentially added to lateral side of existing sarcomeres when the Activin signaling is low or absent. In addition to positively regulating InR/dTOR pathway, the Activin pathway also promotes the expression of putative factors that shift the mode of MHC addition toward forming new sarcomeres and also vertical expansion of existing sarcomeres. Mammalian homologous sites of phosphorylation are presented in parentheses.

## Acknowledgements

We thank Graeme W. Davis and Ernst Hafen for providing *Pdk1* mutants. We are grateful to the Bloomington *Drosophila* Stock Center for providing fly lines and Developmental Studies Hybridoma Bank for α-Actn antibody. We also thank Aidan Peterson, Thomas Neufeld and Hiroshi Nakato for comments on the manuscript. This work was supported by a National Institutes of Health grant (1R35 GM-118029) to M.B.O.

