## Supplemental figures for "*Drosophila* Activin signaling promotes muscle growth through InR/dTORC1 dependent and independent processes"

**Supplementary Figures**

**Fig. S1.** (A) The specificity of the anti-pdSmad2 antibody was examined by immunoblot analysis. The absence of corresponding bands of pdSmad2 in *babo* and *dSmad2* mutants verifies the specificity of the antibody. The absence of pdSmad2 band in *babo* mutants also indicated that dSmad2 phosphorylation is exclusively canonical in larval body wall tissue. (B) As in muscle 6, both the Z-disc number and muscle width are decreased in muscle 7 of *babo* and *dSmad2* mutants. (C) A representative image showing how the Z-discs are detected and intervals are measured from $\alpha$-Actn staining by PeakFinder macro of ImageJ software. (D) Sarcomere size assessed by Z-disc interval is decreased in *babo* but not in *dSmad2* mutant. (E) Sequence alignment of $myo_{CR2}$ mutant line with wild-type. $myo_{CR2}$ has a lesion with one base pair deletion in the target sequence. (F) Number of nucleus from muscle 6 of abdominal segment 2 and 3. The *babo* mutant shows a similar number of nucleus as $w_{1118}$ which we used as a wild type in this study, whereas *dSmad2* mutant exhibits an increased nucleus number compared to $w_{1118}$ (red asterisks). When compared to *yw*, all genotypes including $w_{1118}$ are found to have a smaller number of nucleus (black asterisks) except in the abdominal segment 3 of *dSmad2* mutant. Values are mean ± SEM. $^{**}p<0.01$ and $^{***}p<0.001$ from one-way ANOVA followed by Dunnett's test in which each genotype was compared to *wt*.

**Fig. S2.** (A) Quantification of transcripts level of sarcomeric proteins in *wt* as well as in *babo* and *dSmad2* mutants by qPCR. Transcription of *Mhc* is not significantly altered while *Actn* expression is up-regulated by *babo* and *dSmad2* mutations. (B) Quantification of transcripts level of Zasps in *wt* as well as in *babo* and *dSmad2* mutants by qPCR. Transcription of *Zasp52* is not significantly altered while *Zasp66* expression is down-regulated in *babo* and *dSmad2* mutants. (C) Representative immunoblot image and quantification of MHC. Co-expressed *raptorRNAi* suppressed the hyper production of MHC caused by *babo*$^{CA}$. Values are mean ± SEM. *p<0.05, **p<0.01 and ***p<0.001 from one-way ANOVA followed by Dunnett's test in which each genotype was compared to *wt* (A and B), *Mef2-Gal4/+* control (C). Additionally, an unpaired t-test was performed in C as indicated by lines. ##p<0.01 from unpaired t-test.

**Fig. S3. Reproduction of the key results using *Mhc-Gal4* driver.** (A) Representative immunoblot images of MHC and pAKT1. Overexpression of *dSmad2* transgene using *Mhc-Gal4* driver in *dSmad2* mutant background restores the altered levels of MHC and pAKT1. (B) Representative muscle images stained with Actn antibody. Overexpressing *dSmad2* transgene using *Mhc-Gal4* driver rescues the reduced size of *dSmad2* muscle. Scale bar equals 50 µm. (B') Quantification of muscle size by counting Z-discs. *Mhc-Gal4*-driven expression of *dSmad2* transgene rescues the decreased Z-disc number of *dSmad2* muscle. (C) Overexpressing *babo*$_{CA}$ using *Mhc-Gal4* driver greatly increases the Z-disc number. Values are mean ± SEM. ###p<0.001 from unpaired t-test.

**Fig. S4. Decoupling between myonuclei and muscle sizes.** (A) Representative images of Myc and DPAI staining on *wt*, *dSmad2*$^{F4}$*/Y;Mef2>+* and *dSmad2*$^{F4}$*/Y;Mef2>dSmad2* muscles. Muscular expression of *Myc* increased the myonuclei size but failed to rescue the size of *dSmad2* muscle. (B) Quantification of muscle and myonuclei sizes. (C) The ratio of average myonuclei size to muscle surface area is not altered in *dSmad2* muscle while it is greatly increased by *Myc* overexpression. Values are mean ± SEM. ***p<0.001 from one-way ANOVA followed by Dunnett's test in which each genotype was compared to *wt*. Additionally, unpaired t-tests were performed in as indicated by lines. ns: not significant, ##p<0.01 from unpaired t-test.
